# Elucidating role of long non-coding RNAs of *Tamarindus indica* Linn. in post-transcriptional gene regulation

**DOI:** 10.1101/2024.04.03.587866

**Authors:** Moumita Roy Chowdhury, Aman Kumar, Alfred Besra, Jolly Basak

**Affiliations:** Department of Biotechnology, Indian Institute of Technology Kharagpur, Kharagpur-721302, India; Department of Biotechnology, Koneru Lakshmaiah Education Foundation, Guntur, 522302, Andhra Pradesh, India; Department of Biotechnology, Visva-Bharati University, Santiniketan-731235, India; Department of Biotechnology, St. Xavier college, Pathalkudwa, Nayatoli, Ranchi, Jharkhand-834001, India

**Keywords:** *Tamarindus indica* Linn., lncRNA, CDS, ORFs, motifs, TRs, gene regulation, qRT-PCR

## Abstract

*Tamarindus indica*, commonly known as tamarind, is a rich source of carbohydrates, proteins, lipids, fatty acids, vitamins, minerals and different bioactive compounds. It is well established that long non-coding RNAs (lncRNAs) play important role in transcriptional, post-transcriptional and epigenetic regulation. In spite of the tamarind genome information available, handful of studies have been done on its non-coding genome. In this study, 320 tamarind lncRNAs have been predicted by computational methods. Along with the experimental validation of seven randomly chosen lncRNAs, functional analysis of the predicted lncRNAs along with their targets elucidated their roles in various biological pathways. Sequence analysis of these predicted lncRNAs reveals the presence of different motifs and TRs. Our analysis provides information about the non-coding genome of tamarind and their involvement in gene regulation, which may be used to gain more knowledge about the medicinal properties of tamarind.

## Introduction

Tamarind or *Tamarindus indica* Linn. is one of the most important cultivar of fabaceae family and is widely cultivated in Africa and parts of southern Asia (1). Tamarind is a rich source of carbohydrate, protein, lipid and other nutrients like vitamins (thiamine, riboflavin, niacin and Vitamin C) and minerals (Ca, Mg, Fe, K, Cd, Mn, p, Cr and Zn) (2,3). Tamarind seeds are also rich source of different fatty acids like linoleic, oleic and palmitic acids (4). Various bioactive compounds like alkaloids, flavonoids, steroids, tannins, saponins, phenols, glycosides are also present in the fruit pulp (5,6). Due to the presence of all these important bioactive compounds, over the years, tamarind has been considered as a medicinal plant. Recent studies has proven that *T. indica* has anti-inflammatory and analgesic properties which may be the justification of using tamarind to treat inflammatory diseases like arthritis and body pain (5,7). Presence of alkaloids, flavonoids, steroids are might be the reasons for these anti-inflammatory and analgesic properties (5). It has been found that tamarind has anti-hyperglycaemic activity which may be attributed to glycosides, saponins and anthraquinones present in the fruit pulp (6). Methanol and acetone extracts of tamarind have shown potent antimicrobial activity against *Klebsiella pneumonia*, while the concentrated extracts showed significant antimicrobial activity against *Salmonella paratyphi, Bacillus subtilis, Salmonella typhi*, and *Staphylococcus aureus* (8). Aqueous-ethanol extract of tamarind have shown protective effects against nephrotoxic effects of gentamicin (9). It has been seen in recent study that ethyl acetate extracts from aqueous phase of tamarind pulp contain ellagic acid, chlorogenic acids, caffeic acid and flavons which are responsible for the antioxidant activities of tamarind (10).

Non-coding RNAs are found to be involved in various regulatory processes along with the maintenance and segregation of chromosomes (11). Long non-coding RNAs (lncRNAs), transcribed by RNA polymerase II or III, are transcripts longer than 200 nt bases with an open reading frame (ORF) of less than 100 amino acid (12). Previously lncRNAs were considered as transcriptional noises, but later it has been found that lncRNAs are key players of numerous gene regulation processes including transcriptional, post-transcriptional and epigenetic regulation (13). In plants, lncRNAs are involved in growth, reproduction, development, cell differentiation, chromatin modification, protein re-localization, phosphate homeostasis, and responses to biotic and abiotic stresses (14–16). Plant lncRNAs can be classified based on their genomic location and functions (17). lincRNAs, transcribed from the intergenic regions, regulate gene expression epigenetically and also control protein levels (18). Intronic lncRNAs, transcribed from intronic regions are found to be involved in epigenetic repression (19). Sense and anti-sense lncRNAs are transcribed from the sense and anti-sense strand of protein coding genes and anti-sense lncRNAs are wide spread in plants and well-studied than sense-lncRNAs (20,21). *Cis*-acting lncRNAs can regulate genes transcribed from the same chromosomal locus (22). On the other hand *trans*-acting lncRNAs can relieve the suppression on mRNAs by binding to the miRNA and by acting like a target-mimic for the specific miRNA (23).

In comparison to small non-coding RNAs, lncRNAs are less explored and particularly scanty information is available about plant lncRNAs. Large number of plant lncRNAs can be identified and characterized through the blending of genome-wide computational and experimental approaches (17,20,24–28). Since no lncRNAs have been reported in tamarind so far and lncRNAs can play a major role in the medicinal properties of tamarind, we have identified tamarind lncRNAs and studied their possible roles in this aspect. lncRNAs are predicted in this study using computational methods where coding potentiality and ORF length are considered as significant parameters. In this study, 320 lncRNAs along with their sequence analysis and targets on mRNAs are predicted. We have also experimentally validated randomly chosen seven lncRNAs in this study.

## Materials and methods

### Computational prediction of lncRNAs and their characterization

Coding DNA sequences (CDSs) of *T. indica* with length more than 200 nt were first selected. EMBOSS getorf was used to get the ORFs with length lesser than 100 amino acids (29). Coding Potential Calculator 2.0 (CPC2) (30) and Coding Potential Assessment Tool (CPAT) (31) were used to check coding potentiality of these sequences. BLASTX was performed with SWISS-PROT database as subject and the remaining non-coding sequences as query and the with e-value cut-off of 0.001. Sequences with 40% resemblance or more got removed and rest of the non-coding sequences were considered as lncRNAs. The flowchart of predicting lncRNAs is shown in Fig. 1. Presence of the motifs in the lncRNAs were identified by using DREME (32), and presence of the tandem-repeats (TR) were identified by using Tandem Repeats Finder (33).

**Figure 1.**
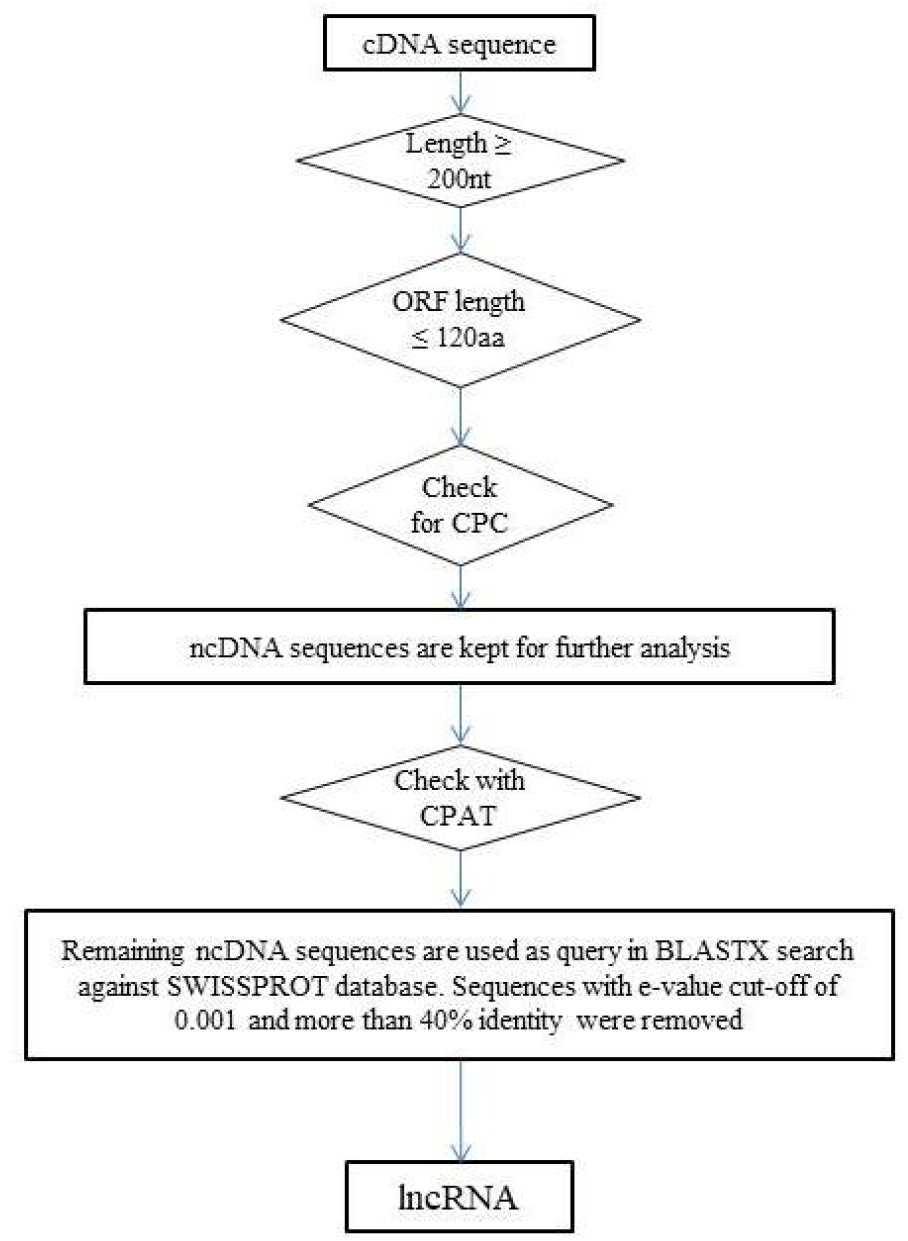
Schematic diagram of the prediction method of lncRNAs. cDNA sequences of *B. oleracea* were downloaded from NCBI. lncRNAs are predicted after filtering out all the criteria discussed in the materials and methods section.

### Prediction of lncRNA targets on mRNAs and functional annotation of the target

In order to predict the targets of lncRNAs on mRNAs LncTar (34) has been used. LncTar is based on calculation of approximate binding free energy, deltaG (dG), of each pairing which indicate the stability of complementarity between lncRNA and mRNA. Predicted mRNA targets were analyzed for their functional annotation.

### Plant materials for validation of the lncRNAs

Tamarind seeds were surface sterilized by using 0.1% HgCl_2_ for 2-5□minutes and then washed with distilled water. Seeds were sowed in sterile soilrite and germinated in dark. Growth of seedlings was maintained for ten days in plant growth chamber with a photoperiod of 16/8 hour with 70–80% relative humidity at 24°C.

### Total RNA Isolation

Total RNA was isolated by using Macherey-Nagel nucleospin plant RNA isolation kit from the leaves of ten days old tamarind seedlings. Isolated total RNA was quantified by spectrophotometer (model: Eppendorf).

### cDNA synthesis and primer design

First-strand cDNA synthesis kit (Thermo) was used to transcribe cDNA from the isolated RNA following the manufacturer’s instructions. Primers (forward and reverse) were designed using Primer3 (https://primer3.org/) and are listed in table 1.

**Table 1:**
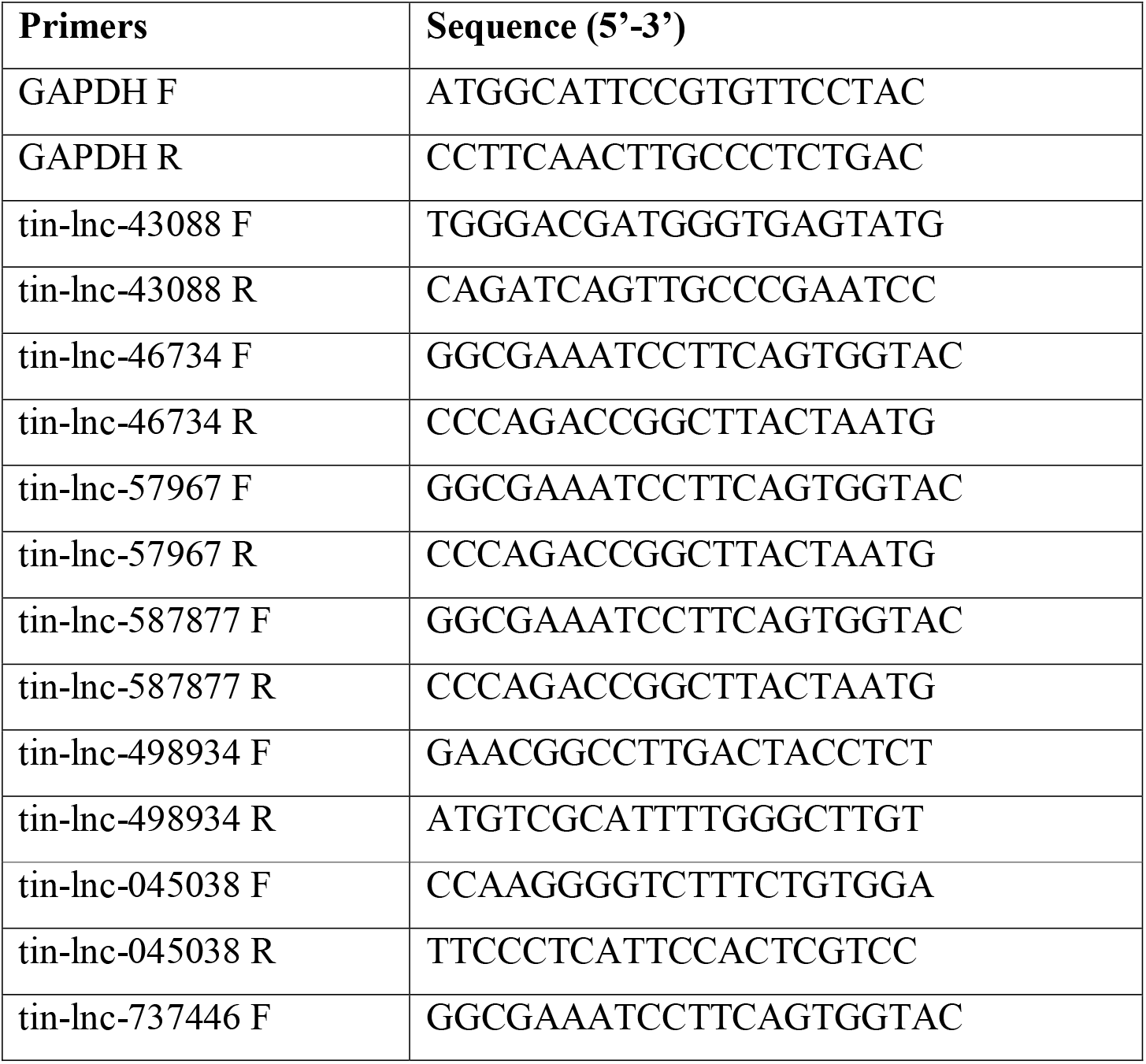
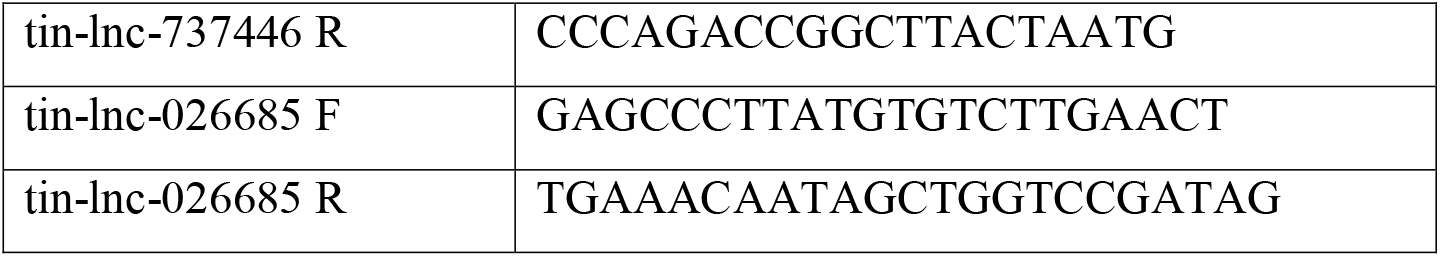
Primers used for RT-qPCR assay.

### Quantitative Real Time Polymerase Chain Reaction (qRT-PCR)

For qRT-PCR, Bio-Rad CFX96 RT-PCR system along with SYBR green supermix (Bio-Rad iQ) were used. Each reaction mixture (10 μL) contained cDNA, SYBR green supermix and 300 nM of forward and reverse primers. The PCR cycle protocol was followed as mentioned in Roy Chowdhury et al. (2021). Specificity of the amplicons were verified by melting curve analysis. Each reaction was executed in triplicates and confirmed by occurrence of a single peak of the melt curve. Glyceraldehyde 3-phosphate dehydrogenase (GAPDH) was used as endogenous control.

### Statistical analysis

Each data has been tested separately for statistical analysis. Data is represented as mean ± standard error of means (SEM). Student’s t test was performed for identifying significant changes by taking p < 0.05 as threshold. All the tests have been performed on independent biological triplicates.

## Results

### Computational prediction of lncRNAs and their characterization

In this study we have predicted 320 lncRNAs (supplementary table S1). Most of the lncRNAs are greater than 400nt in length. Length of the remaining lncRNAs ranges between 200-400 nt. Length distribution of the lncRNAs is shown in Fig. 2. We have found five different motifs in the computationally predicted lncRNAs which are involved in different biological processes, molecular functions and being part of cellular components. Detailed description of the motifs and their functional significances are described in table 2. In this study we have predicted TRs and we found that seven lncRNAs are having simple repeats, LINE/CR1 repeats as well as low complexity regions (table 3).

**Figure 2.**
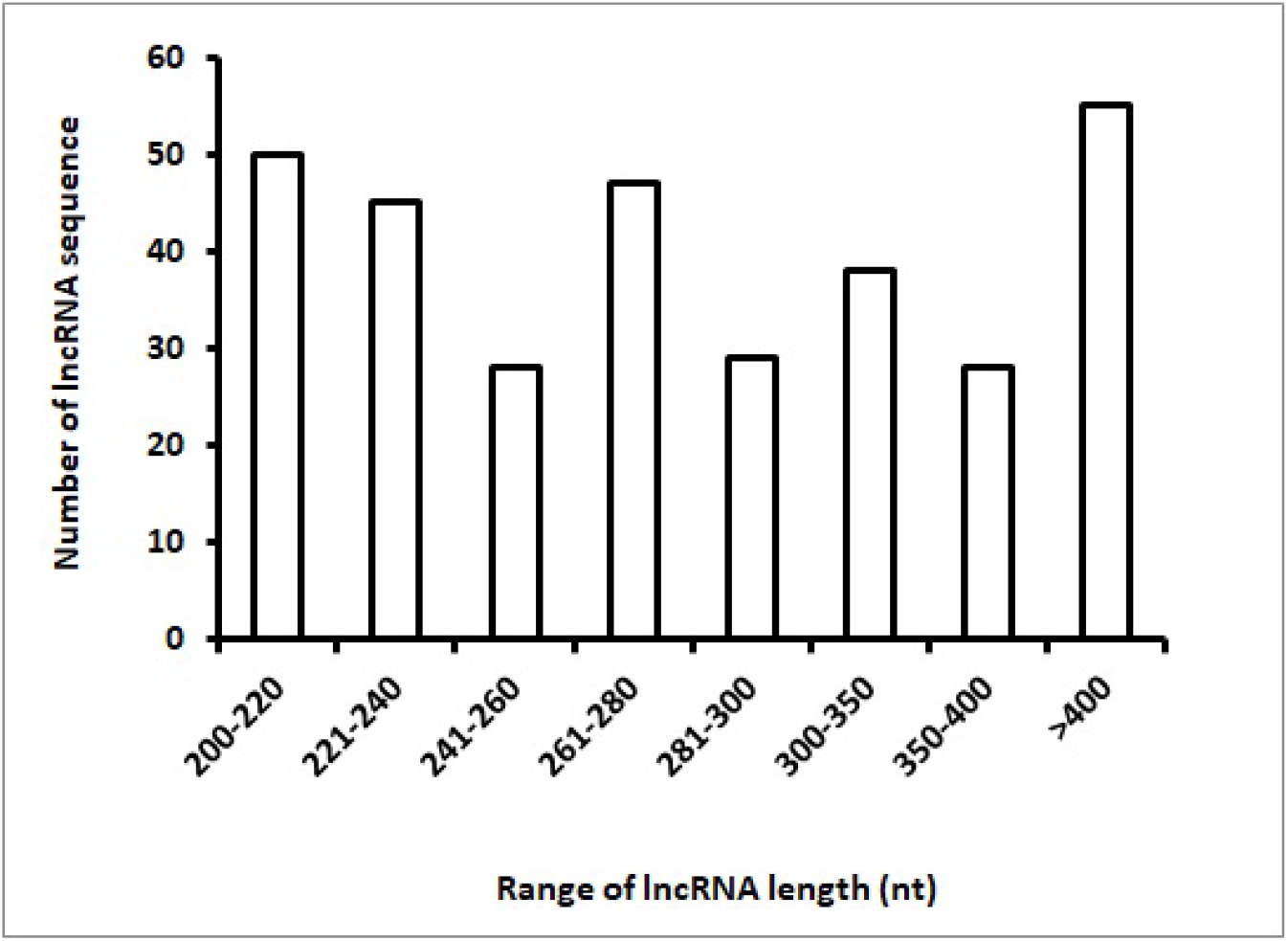
Distribution of lncRNA based on their length.

**Table 2:**
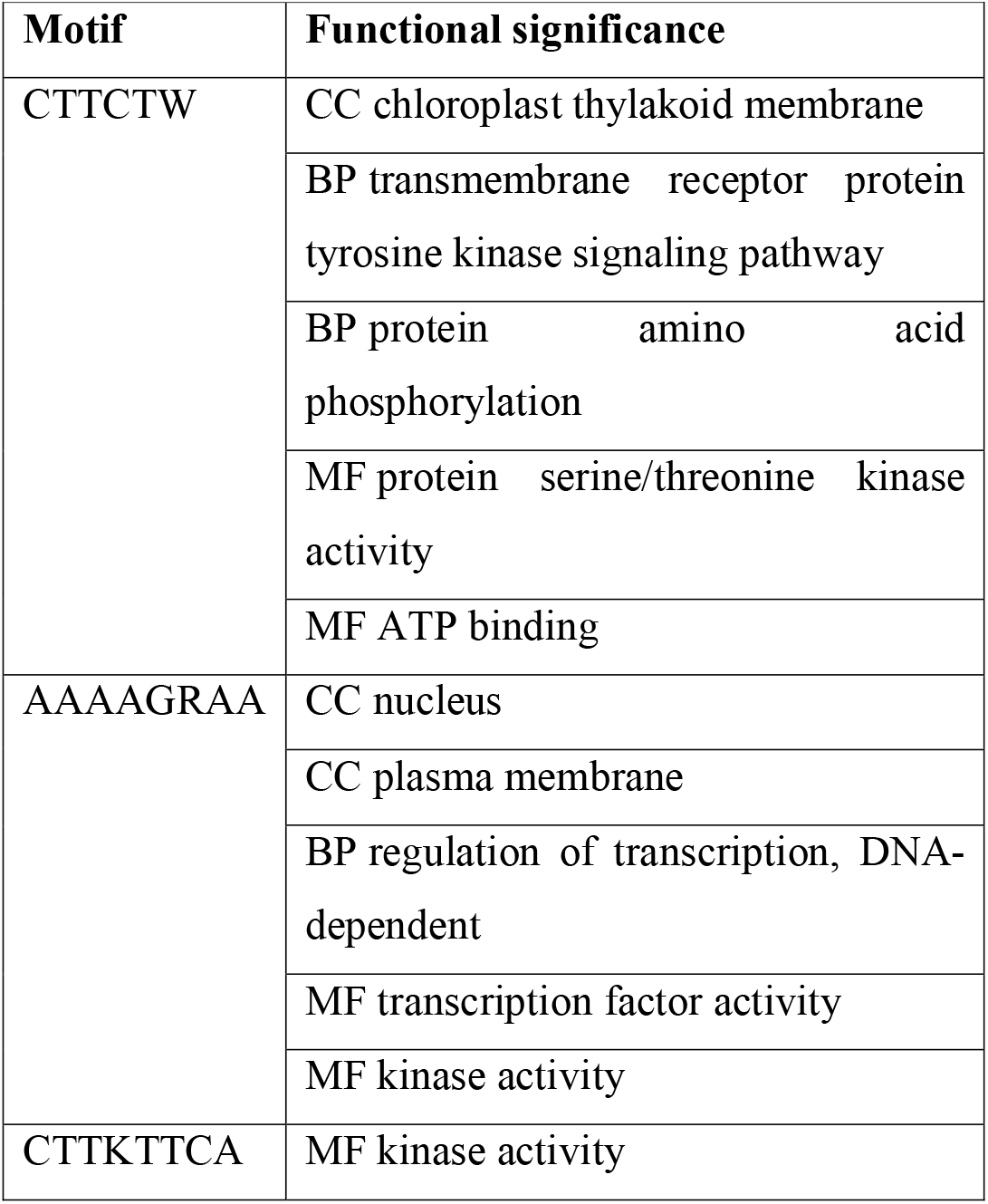
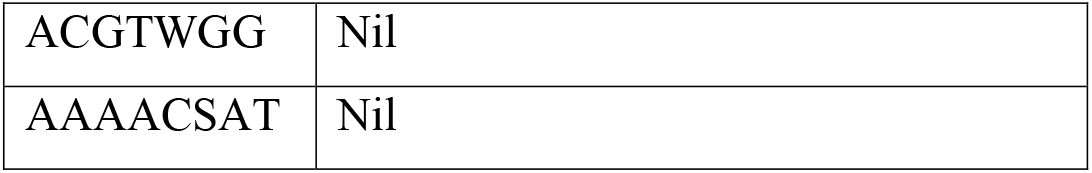
Motifs and their functional significance.

**Table 3:** Repetitive elements in the lncRNAs.

| **Repeat class/family** | **lncRNAs** |
| --- | --- |
| Simple repeats | tin-lnc-0000023c, tin-lnc-0000026a, tin-lnc-0000036d |
| Low complexity regions | tin-lnc-026685f |
| LINE/CR1 | tin-lnc-0000029f |

### Prediction of lncRNA targets on mRNAs and functional annotation of the targets

Among the lncRNAs, eight lncRNAs are found to have targets on five protein coding genes. Seven lncRNAs namely, tin-lnc-026685c, tin-lnc-026685l, tin-lnc-0000037, tin-lnc-0000034, tin-lnc-0000024e, tin-lnc-000009f and tin-lnc-000009i are found to target a single gene UDP-sulfoquinovose synthase (SQD1) protein coding gene while tin-lnc-498934 targets 4 different FLO/LFY-like protein coding genes (A4, A5, A6 and A7). UDP-sulfoquinovose synthase is involved in nucleotide sugars metabolism and glycerolipid metabolism. Floricaula/leafy-like transcription factor is involved in regulation of transcription, DNA binding.

### Validation of the predicted lncRNAs

From the computationally predicted lncRNAs, seven lncRNAs are randomly selected and are experimentally validated. Relative quantity of lncRNA expression is gained by 2^-ΔC^T, where ΔC_T_ = (C_T_ lncRNA-C_T_ GAPDH). Amongst the seven lncRNAs, C_T_ value of tin-lnc-587877 is the lowest (17.385), followed by tin-lnc-737446 (C_T_ = 18.8), tin-lnc-57967 (C_T_ = 19.26) and tin-lnc-498934 (C_T_ = 19.49). For the remaining three lncRNAs, C_T_ values of two lncRNAs namely tin-lnc-045038 (C_T_ = 23.6) and tin-lnc-026685 (C_T_ = 25.4) are in medium range while tin-lnc-43088 (C_T_ = 30.6) has the highest C_T_ value. Expression profiles of the seven lncRNAs from qRT-PCR analysis are shown in Fig. 3.

**Figure 3.**
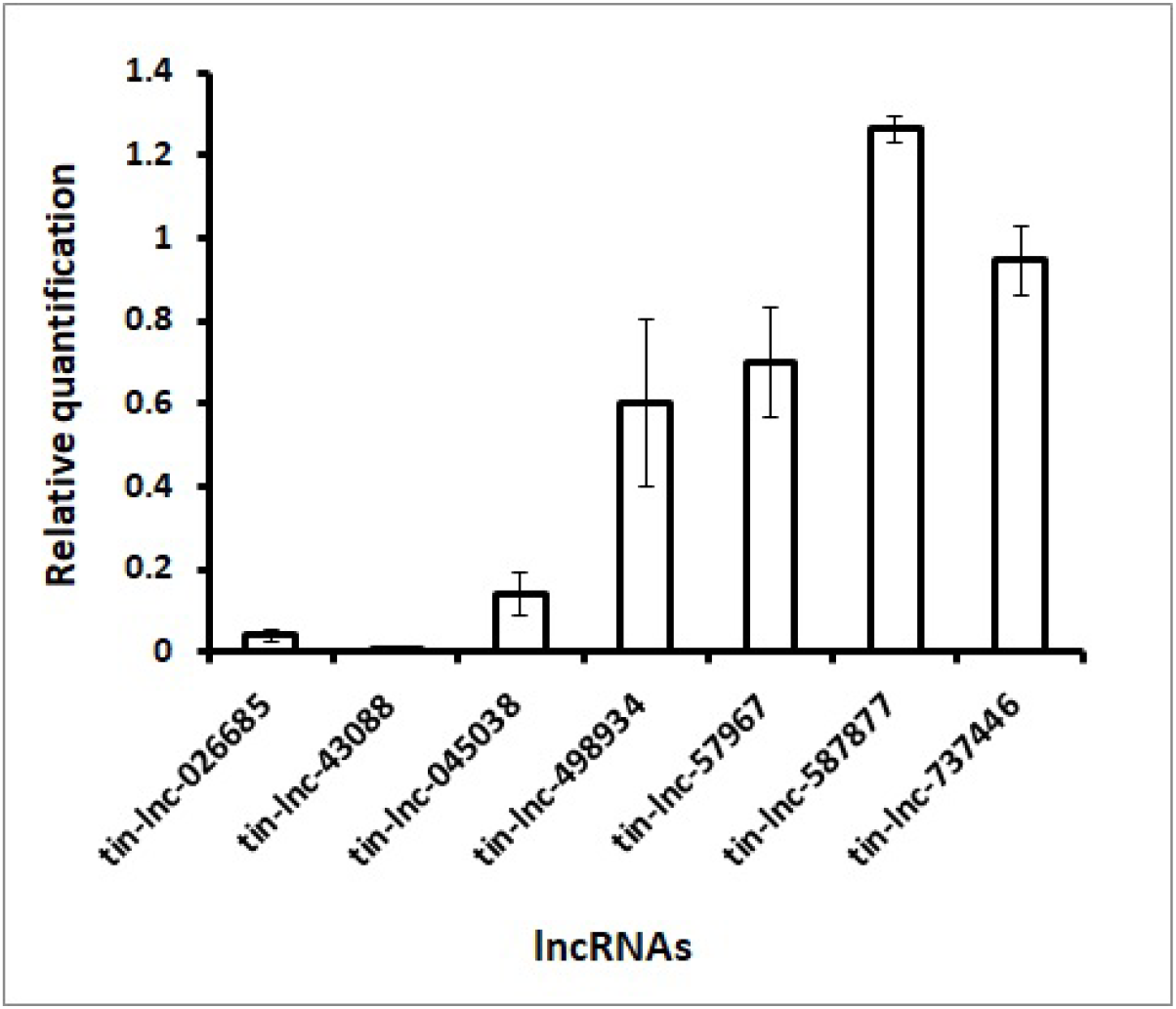
Expression profile of 10 lncRNAs from qRT-PCR analysis. Relative quantification was measured by 2^-ΔCT^ where ΔC_T_ = (C_T_ lncRNA-C_T_ GAPDH).

## Discussions

Presence of very less information on tamarind lncRNAs has prompted us to perform this study in order to gather more information on tamarind lncRNAs. Being a rich source of nutrients and secondary metabolites, tamarind possesses immense importance in human diet (2,3). Although there is information available about the coding genome of tamarind, very less information is available on non-coding genome, especially on lncRNAs. In recent years, it has been found that plant lncRNAs play important roles in various biological processes like growth, development, cell differentiation, protein-re-localization, response to different biotic and abiotic stresses, phosphate homeostasis and chromatin modification (14–16). Through this study, we have computationally predicted new lncRNAs and also tried to find out their functional significances. Besides predicting novel lncRNAs of tamarind, we have also identified the involvement of these novel lncRNAs in different biological pathways by predicting their target mRNAs followed by their functional annotation. In this study, we have seen single lncRNA targeting multiple protein coding genes as well as multiple lncRNAs targeting single protein coding gene. We found SQD1, an enzyme involved in biosynthesis of sulfoquinovosyl headgroup of plant sulfolipids and catalyzing the transfer of SO_3_^−^ to UDP-glucose (36), is targeted by seven different lncRNAs. On the other hand, tin-lnc-498934 targets four different FLO/LFY-like protein coding genes (A4, A5, A6 and A7), which regulate first cell division after the zygote formation (37). Amongst the five motifs present in these lncRNAs, functional annotation is carried out for three motifs in this study. Motif “CTTCTW” is involved in biological processes like trans-membrane receptor protein tyrosine kinase signaling pathway and protein amino acid phosphorylation. It also takes part in molecular functions like protein serine/threonine kinase activity and ATP binding. Motif “AAAAGRAA” is involved in biological process like regulation of transcription along with transcription factor activity like molecular functions. Motif “CTTKTTCA” is involved in kinase activity. For the remaining two motifs, functional annotation did not predict any function, neither biological process nor molecular function. We have also predicted TRs that are important driving factors on the emergence of lncRNAs (38). Three lncRNAs namely, tin-lnc-0000023c, tin-lnc-0000026a, tin-lnc-0000036d consist of simple repeats “ATT”, “GAAT”, “CTTTTT” respectively. Two lncRNAs namely tin-lnc-026685f and tin-lnc-0000029f consist of low complexity regions and LINE/CR1 repeats respectively. Randomly chosen seven lncRNAs were experimentally validated. tin-lnc-587877 showed the highest expression. Other highly expressed lncRNAs are tin-lnc-737446, tin-lnc-57967 and tin-lnc-498934. Two moderately expressed lncRNAs are tin-lnc-045038 and tin-lnc-026685, whereas tin-lnc-43088 showed lowest expression.

## Conclusion

We have identified lncRNAs from the coding DNA sequence of *T. indica* and experimentally validated randomly chosen seven lncRNAs. Further, we have predicted five targets for four lncRNAs. We have identified the motifs and the TRs, along with functional prediction of the motifs. We strongly believe that our study will definitely contribute to the current knowledge of the tamarind non-coding genome which is still in its infancy. Prediction of lncRNA, their targets along with involvement of the targets in diverse biological pathways will provide more insights into the non-coding genome of tamarind, as well as may shed light to many unknown functions related to medicinal and therapeutical properties of tamarind.

## LIST OF ABBREVIATIONS

TRs: Tandem Repeats
lncRNAs: long non-coding RNAs
ORF: open reading frames
CDS: Coding DNA sequence

## ETHICS APPROVAL AND CONSENT TO PARTICIPATE

Not applicable.

## AUTHORS’ CONTRIBUTIONS

MRC, AK and AB performed the study. MRC wrote the manuscript. JB participated in its design and coordination.

## ACKNOWLEDGEMENT

MRC acknowledge IIT Kharagpur for research fellowship and Visva-Bharati university for providing the research facilities for wet-lab experiments. JB acknowledge CSIR for their support.

## DATA AVAILABILITY STATEMENT

All data generated or analyzed during this study are included in its supplementary information files.

